# Spatiotemporal information is differentially conveyed by hippocampal projections to the anterior olfactory nucleus during episodic-like odour memory

**DOI:** 10.1101/230185

**Authors:** Afif J. Aqrabawi, Jun chul Kim

**Author notes:** Correspondence and requests for materials should be addressed to J.C.K.

## Abstract

The hippocampus is essential for representing spatiotemporal context and associating it with the sensory details of daily life to form episodic memories. However, the neural circuit underlying this process remains poorly understood. We selectively inhibited hippocampal projections to the anterior olfactory nucleus (AON) during behavioural tests of contextually-cued odour recall. We found that inhibition of intermediate HPC (iHPC)-lateral AON (lAON) pathway impaired spatial odour memory while inhibition of ventral HPC (vHPC)-medial AON (mAON) pathway disrupted both spatial and temporal odour memory. Our results indicate that the spatial and temporal information of episodic-like odour memory is conveyed by topographically distinct hippocampal-AON pathways.

---

What happened, when, and where? The ability to readily integrate elements of a unique event into a single representation is a fundamental property of episodic memory^1^. Lesion and recording studies in humans and nonhuman animals have highlighted a central role of the hippocampus (HPC) in mediating episodic memory^2, 3^. The spatial and temporal context of an event are first encoded within the HPC as the collective activity of place and time cells^4, 5^. Contextual information later serves as a potent retrieval cue, bringing about the rich multisensory details of the original experience^6, 7^. An emerging theory holds that the HPC conducts this retrieval process by reinstating patterns of cortical activity observed during learning^8, 9^. Yet, it is not known how hippocampal transmission of contextual information can reproduce the sensory aspects of episodic memory.

Olfaction is considered the most evolutionarily ancient sense as evidenced by the direct anatomical connections between the olfactory cortex and the limbic system^10^. In particular, hippocampal projections to the olfactory cortex offer a unique experimental model for understanding the context-driven recollection of previously encountered sensory stimuli. Recently, we revealed a dense and topographically organized projection from the dorsoventral extent of the HPC to the anterior olfactory nucleus (AON) (Fig. 1a; Aqrabawi and Kim, 2017). The AON receives unidirectional, monosynaptic inputs from the CA1, in contrast to other primary sensory areas that receive hippocampal inputs indirectly via adjacent medial temporal lobe structures, underlining further the intimate relationship between olfaction and memory^11^.

**Figure 1.**
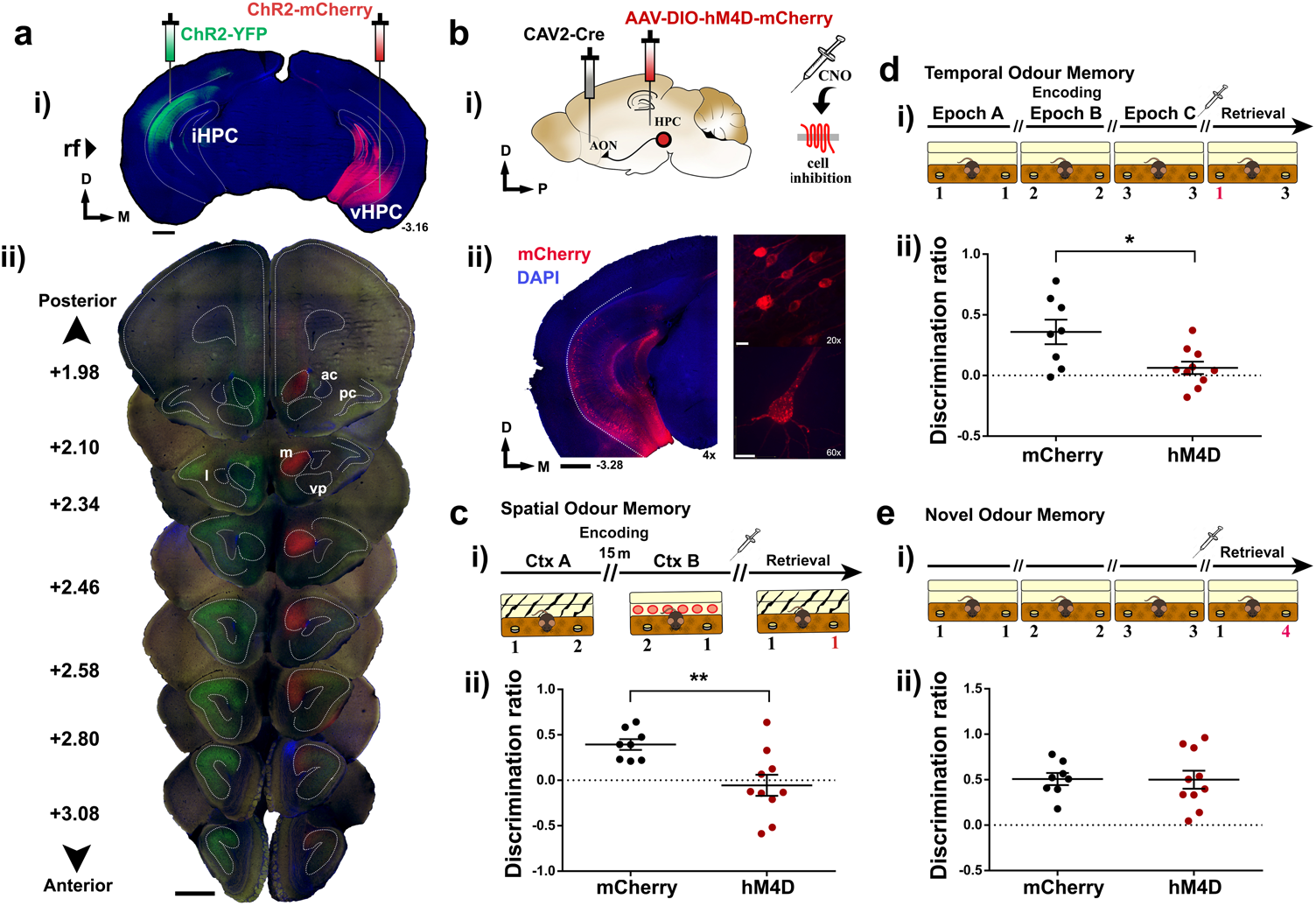
Inhibition of AON-projecting hippocampal neurons impairs retrieval of context-associated memory for previously encountered odours. (**a**) i-ii) Coronal section depicting site of AAV-ChR2-YFP and -mCherry injections in the hippocampus (i) and the resulting innervation pattern at the AON (ii). Coordinates mark anteroposterior position from bregma. Scale bars represent 1 mm; ac= anterior commissure; pc= piriform cortex; l= lateral AON; m= medial AON; vp= ventroposterior AON. (**b**) i) Schematic diagram depicting experimental approach. CAV2-Cre was infused into the AON whereas Cre-responsive AAV-hM4D-mCherry was injected in the hippocampus. ii) Representative sections depicting AON-projecting HPC neurons expressing hM4D-mCherry. Magnification is indicated on the bottom right of each panel. Scale bars represent 1 mm (black) or 10 μm (white). (**c**) i) The olfactory spatial memory test paradigm. ii) CNO-treated hM4D mice investigated the familiar and novel odour location to a largely equal extent, indicative of impaired spatial odour memory (Independent-samples t-test, t(i6)=3.194, \*\**P*<0.01). (**d**) i) The olfactory temporal order memory test paradigm. ii) CNO-treated hM4D mice are impaired in memory for the temporal occurrence of encountered odours (Independent-samples t-test, t_(16)_=2.795, \**P*<0.05). (**e**) i-ii) Both groups showed normal performance in a context-independent novel odour recognition test (Independent-samples t-test, t_(16)_=0.05644, ns, *P*=0.9557). For all behavioural tests, CNO was injected 15 minutes prior to the retrieval phase. The odour in the novel spatiotemporal configuration is numbered in red. Positive discrimination ratios indicate preference for the novel odour-context configuration. mCherry control group: n=8, hM4D group: n=10. Data are presented as mean ± s.e.m.

The AON is an ideal site of convergence for olfactory and contextual information given its anatomical position as the initial recipient of input from the olfactory bulb and the largest source of feedback projections within the olfactory cortex^12, 13^. Consistently, it has been shown that hippocampal inputs to the AON can alter olfactory perception and odour-guided behaviours^14^. However, the functional role of the HPC-AON pathway remains unexplored, leaving open fundamental questions regarding olfactory episodic memory. Here, we combine chemogenetic and optogenetic approaches to demonstrate that information regarding the spatial and temporal context of odour memory is delivered by topographically organized hippocampal inputs to the AON.

To manipulate activity in the HPC-AON pathway, we infused the retrogradely propagating canine adenovirus encoding Cre recombinase (CAV2-Cre) into the AON, followed by HPC infusions of an AAV vector carrying a Cre-dependent inhibitory hM4D-mCherry. This allowed us to selectively inhibit HPC cells that project to the AON upon the administration of clozapine-*N*-oxide (CNO). Cre-mediated viral mCherry expression was observed throughout the ventral two thirds of the hippocampal CA1 (Fig. 1b). Three weeks after viral infusion, mice underwent behavioral tests to evaluate memory for the associations between odours and the spatiotemporal context in which they occurred. Assessing the retrieval of episodic odour information was made possible by capitalizing on the innate tendency of mice to preferentially investigate novel stimuli^15^. Thus, mice can behaviourally express correct memory by investigating odours paired with a novel position in space, or temporal sequence, more so than familiar configurations.

We first tested the ability to remember ‘where’ specific odours occurred in context. In two encoding phases of a spatial odour memory test, mice were presented with two different odours (odour 1 and 2) placed on opposite ends of a distinct context (A) for 5 min (Fig. 1ci). Mice were then placed in a separate context (B) where the same odours were positioned in reversed locations. Each exposure was followed by a 15 min retention delay, the second of which was preceded by a CNO injection. In the subsequent retrieval phase, animals were reintroduced to context A wherein two copies of either odour 1 or 2 were presented on both sides. In this paradigm, correct memory expression would drive mice to investigate the odour found at the novel location within the context, as seen in control mice expressing mCherry alone (Fig. 1cii). In contrast, CNO-treated hM4D mice investigated both, otherwise identical odours for a similar proportion of time. These results cannot be explained by differences in total investigation time or distance traveled as both groups displayed similar measures in each (Supplementary Fig. 1).

Next, we examined memory for ‘when’ specific odours were encountered. Mice were presented with a sequence of odours in three successive encoding phases followed by a retrieval phase where two of the previously encountered odours were reintroduced (Fig. 1di). Control mice preferentially investigated the odour encountered earlier in the sequence, yet CNO-treated hM4D mice investigated both odours to a largely equal extent (Fig. 1dii). Importantly, the spatial context was consistent throughout the test where novelty was only conferred by temporal distance between the two odours. Lastly, to examine whether hippocampal function can be extended to memory for odours regardless of context, we conducted a novel odour recognition test where animals were presented with a familiar and previously unexplored odour (Fig. 1ei). Both groups preferentially investigated the novel odour despite CNO administration (Fig. 1eii). Together, these results indicate that AON-projecting HPC cells mediate the retrieval of odour memory only when it is tied to spatiotemporal context.

Topographically organized HPC terminals at the AON may transmit distinct contextual cues depending on where they arise within the HPC. We employed archaerhodopsin (ArchT) to optogenetically inhibit intermediate or ventral HPC terminals that innervate separate AON subregions. One group received bilateral infusions of AAV-CaMKIIa-ArchT-eYFP into the intermediate hippocampus (iHPC) with optic fiber implantations in the lateral AON (lAON) for inhibiting the iHPC-lAON pathway, and another underwent viral infusions into the ventral hippocampus (vHPC) with optic fiber implantations in the medial AON (mAON) for inhibiting the vHPC-mAON pathway (Fig. 2a; Supplementary Fig. 2). A control group given counterbalanced infusions and implantations were prepared using AAV expressing GFP only. Animals were tested in the same behavioural paradigms used for hM4D experiments, except inhibition was mediated by light (532 nm, 12 mW) limited to the retrieval phase.

**Figure 2.**
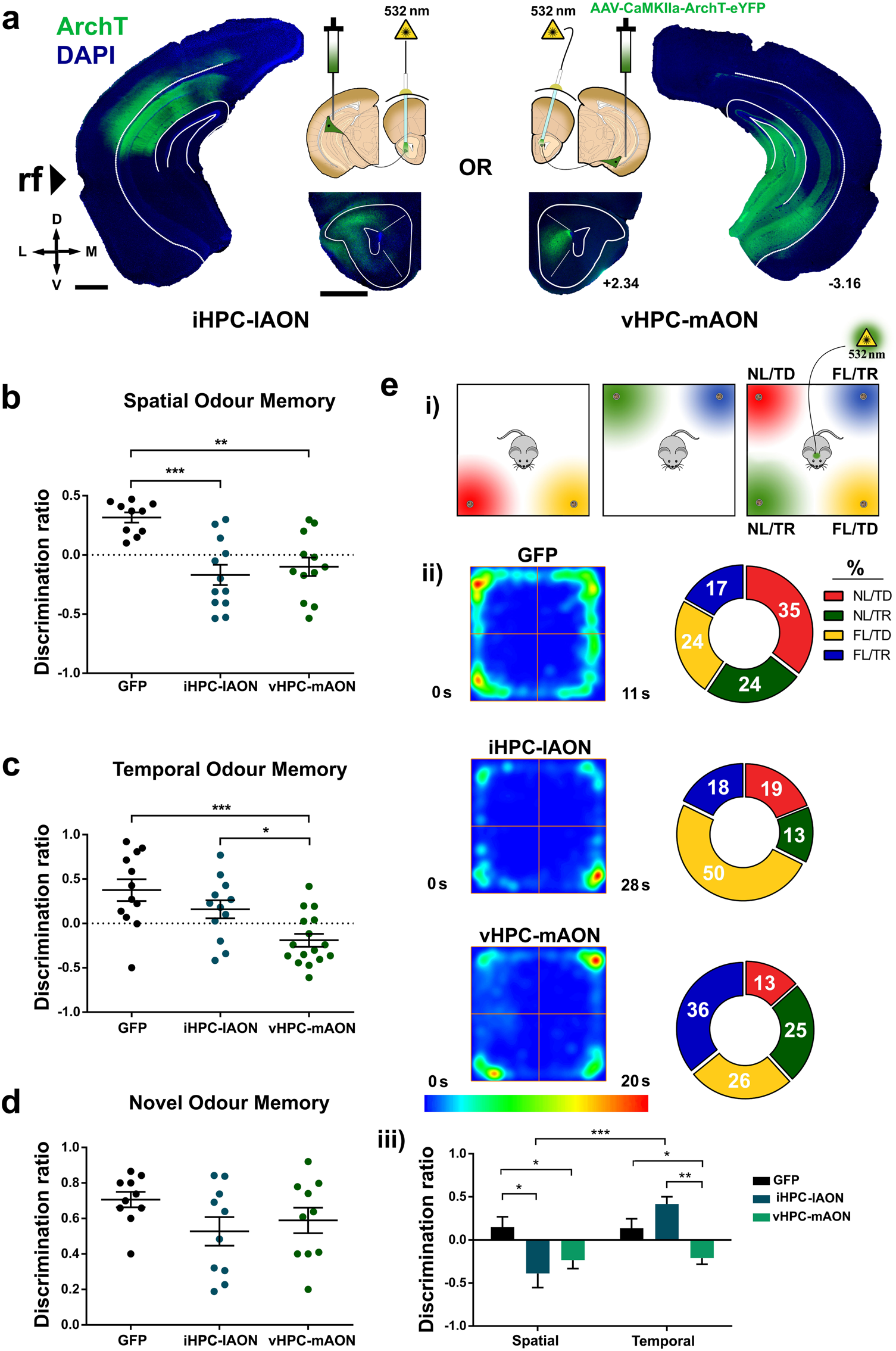
HPC-AON pathways contribute distinct spatiotemporal information during episodic odour memory retrieval. (**a**) Diagram depicting location of ArchT infusions within the HPC and optic fiber implantations above the AON in both experimental groups. iHPC terminals are predominantly found in the lAON whereas vHPC fibers preferentially innervate the mAON. (**b**) Inhibition of both iHPC-lAON and vHPC-mAON pathways impaired spatial odour memory (GFP: n=10; iHPC-lAON: n=12; vHPC-mAON: n=12; one-way ANOVA, F_(231)_=11.65, \*\*\**P*<0.0005). (**c**) Inhibition of the vHPC-mAON, but not iHPC-lAON pathway impaired memory for the temporal order of a sequence of odours (GFP: n=12; iHPC-lAON: n=12; vHPC-mAON: n=16; one-way ANOVA, F_(2,37)_=9.235, \*\*\**P*<0.005). (**d**) HPC-AON pathway is not necessary for context-independent novel odour recognition (all groups: n=10; one-way ANOVA, F_(2,27)_=1.824, ns, *P*=0.1807). (**e**) i) Illustration of the episodic memory test paradigm. ii) Heat maps depicting group average position (left) and pie charts representing the percent investigation time (right) during the retrieval phase of the episodic memory test (all groups: n=12). iii) iHPC-lAON pathway inhibition impaired the spatial, but not temporal component of odour memory, while vHPC-mAON pathway inhibition impaired both components to a similar extent (all groups: n=12; two-way ANOVA treatment-by-memory type interaction F_(2,66)_= 9.320, \*\*\**P*<0.0005; main effect of treatment F_(2,66)_=6.715, \*\**P*<0.005; main effect of memory type F_(1,66)_= 9.620, \*\**P*<0.005). Data are presented as mean ± s.e.m. Scale bars represent 1 mm. Coordinates indicate anteroposterior position from bregma. NL/TD: novel location/temporally distant; FL/TD: familiar location/temporally distant; NL/TR: novel location/temporally recent; FL/TR: familiar location/temporally recent.

Illumination of the iHPC-lAON and vHPC-mAON pathways disrupted memory for ‘where’ odours were encountered as both groups were unable to discriminate odours tied to a spatial location within a specific context (Fig. 2b). In contrast, only vHPC-mAON, but not iHPC-lAON pathway inhibition impaired memory for ‘when’ odours occurred in a temporal sequence (Fig. 2c). Importantly, all groups showed similar levels of novelty preference in a context-independent novel odour recognition test (Fig. 2d). The identified impairments in episodic odour memory retrieval suggest that representations of space and time are differentially distributed across the AON whereby both iHPC and vHPC inputs deliver spatial information, but information regarding the temporal context is supported only by vHPC inputs.

To investigate further the spatiotemporal contributions of hippocampal inputs to the AON, we employed an episodic memory test where recollection of an odour, its spatial location, and temporal occurrence (what-when-where) were tested simultaneously^16, 17^. The test involved two encoding phases and one retrieval phase, each separated by a one hour delay (Fig. 2ei). Light-mediated inhibition was limited to the retrieval phase. During the encoding phases, mice sampled two distinct odours placed at adjacent corners of an open field, and then sampled a new set of odours positioned on the opposite side of the arena. In the retrieval phase, all four odours were presented, each possessing a unique spatiotemporal configuration. Encountered odours were either in a novel location and earlier in temporal distance (NL/TD), familiar in location/temporally distant (FL/TD), novel in location/temporally recent (NL/TR), or familiar in location/temporally recent (FL/TR). The control group displayed a pattern of investigation indicative of their novelty preference such that the greatest proportion of time was spent investigating the odour encountered in the most novel spatiotemporal combination (NL/TD), while investigation time for the remaining odours decreased in a familiarity-dependent manner: (NL/TD) > (FL/TD) ⋍ (NL/TR) > (FL/TR) (Fig. 2eii; Supplementary Fig. 3). The iHPC-lAON group spent the greatest proportion (~50%) of time investigating the odour with the FL/TD configuration while the NL/TR odour was investigated least. Strikingly, inhibition of the vHPC-mAON pathway produced an inverse pattern of investigation to the control group. The odour presented in the FL/TR configuration was investigated most, while the NL/TD odour was investigated least. A separate analysis delineating the spatial and temporal contributions to memory revealed that iHPC-lAON pathway inhibition impaired spatial but not temporal odour associations, while the vHPC-mAON pathway inhibition impaired both components to a similar extent (Fig. 2eiii). Collectively, these findings confirm the differential roles of the iHPC-lAON and vHPC-mAON pathways in conveying spatiotemporal information during olfactory episodic memory retrieval.

**Figure 3.**
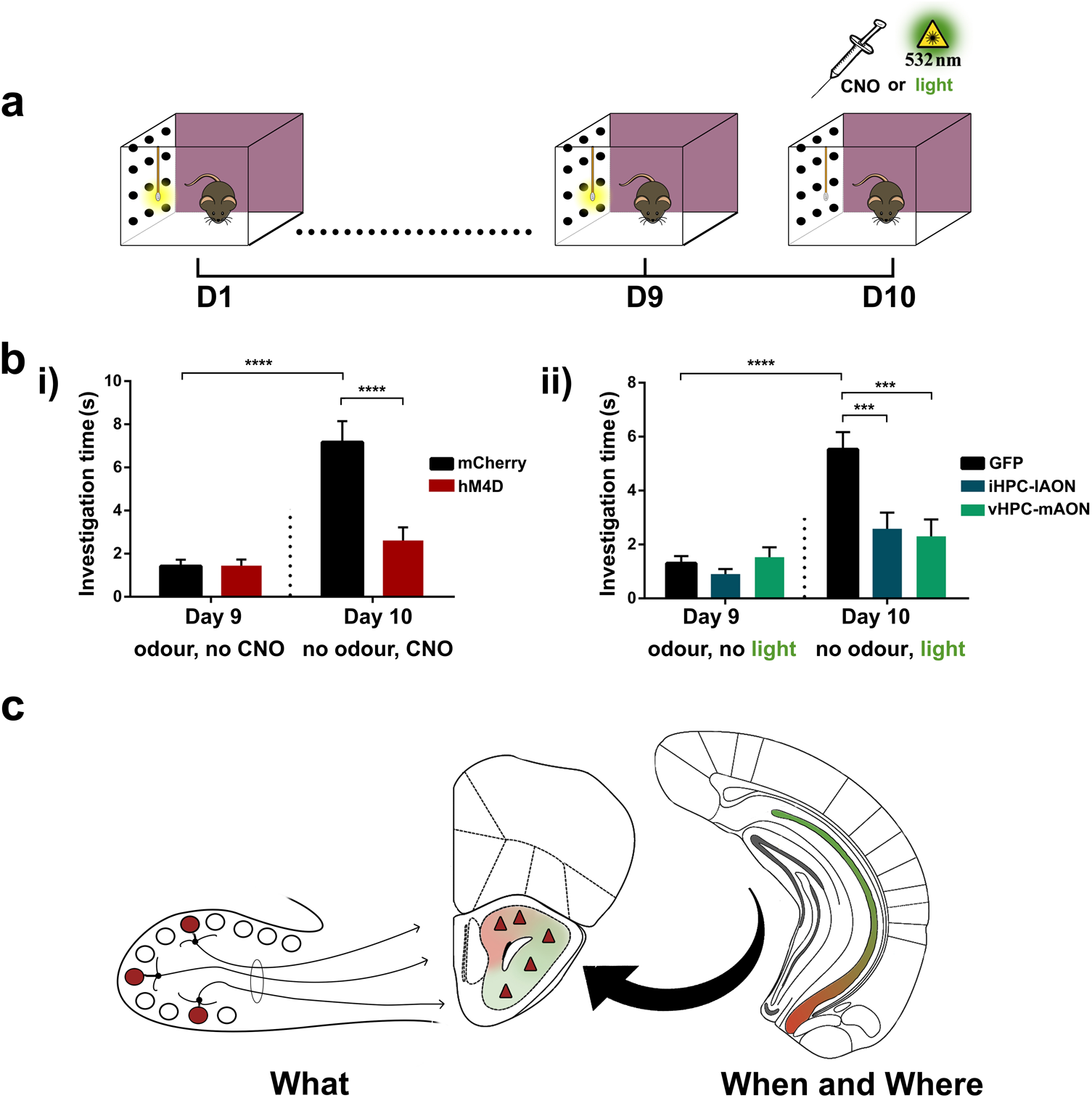
The HPC-AON circuit is necessary for context-driven odour recollection. (**a**) contextually-cued odour recall test paradigm. (**b**) i-ii) Control groups in both experiments showed an increased investigation time between Day 9 and 10. Yet, following hM4D- (i) or ArchT-mediated (ii) inhibition of hippocampal terminals at the AON, mice fail to exhibit this behaviour (hM4D experiment-hM4D-mCherry: n=10, mCherry control: n=8; two-way ANOVA main effect of treatment F_(1,32)_=14.85, \*\*\**P*<0.0005; main effect of Day F_(1,32)_= 34.5, *P*<0.0001; interaction between treatment and Day F_(1,32)_=15.19, \*\*\**P*<0.0005; ArchT experiment-GFP: n=14, iHPC-lAON: n=12, vHPC-mAON: n=12; two-way ANOVA main effect of treatment F_(2,70)_=7.538, \*\**P*<0.005; main effect of Day F_(1,70)_=31.45, \*\*\*\**P*<0.0001; interaction between treatment and Day F_(2,70)_=7.052, \*\**P*< 0.005). Data are presented as mean ± s.e.m. (**c**) A model of episodic odour memory whereby information regarding odour quality and spatiotemporal context merge at the level of the AON producing cellular populations that represent previously encountered odours (what) within the context in which they occurred (when and where).

The ability to explicitly recollect sensory information is a hallmark of episodic memory, particularly when lacking a physical sensory cue^18^. Context alone can drive the activity of primary olfactory regions to form internal representations of odours although the underlying neural circuit is unknown^6, 7^. To this end, we examined whether the HPC-AON pathway could support this function. Mice were allowed to explore a rich spatial context in the presence of a pure odour emitted from a cotton swab tip for 30 min per day over nine consecutive days. On day 10, mice were reintroduced to the context in the absence of the applied odour (Fig. 3a). The mismatch between the odour-paired context and the lack of an emitted scent drives an increase in the investigation time for the cotton swab (Supplementary Video 1). Indeed, control mice exhibited a marked increase in investigation of the cotton swab upon failure to find the expected odour (Fig. 3b). In contrast, hM4D- and ArchT-mediated inhibition of the HPC-AON pathway abolished this behaviour. All groups showed no increase in investigation time when similarly trained but tested in a novel context (Supplementary Fig. 4). Therefore, HPC-AON communication is necessary for mediating the context-driven recollection of odours.

Ultimately our findings support a model of episodic odour memory whereby information regarding odour quality and spatiotemporal context merge at the level of the AON and, as a natural consequence of hebbian synaptic plasticity, produces cellular populations that represent previously encountered odours within the context in which they occurred (Fig. 3c). Such a system maintains the fidelity of the original odour memory and allows access for retrieval of the full trace via partial cues from either olfactory or contextual inputs.

## Methods

### Animals

Male C57BL/6 mice (Charles River Laboratories) were used in all behavioural tests. All mice were 8-10 weeks old at the time of surgery and 12-14 weeks old at the time behavioural testing began. Prior to surgery, mice were group-housed in a temperature-controlled room on a 12 hour light/dark cycle with *ad libitum* access to food and water. Following surgery, mice were individually-housed. A total of 60 mice were distributed into two groups for hM4D experiments (hM4D-mCherry: n=10, mCherry control: n=8) and three groups for ArchT experiments (GFP control: n=14, iHPC-lAON: n=12, vHPC-mAON: n=16). All procedures were performed in accordance with the guidelines of the Canadian Council on Animal Care (CCAC) and the University of Toronto Animal Care Committee.

### Surgical procedures

CAV2-Cre viral vector was purchased from the Plateforme de Vectorologie de Montpellier, AAV2/8-hSyn-FLEX-hM4D-mCherry (hM4D) and AAV2/8-hSyn-DIO-mCherry (mCherry-control) from the vector core at University of North Carolina at Chapel Hill, and AAV2/5-hSyn-hChR2-mCherry (ChR2-mCherry), AAV2/5-hSyn-hChR2-eYFP (ChR2-YFP), AAV2/5-CaMKIIa-ArchT-eYFP (ArchT) and AAV2/8-CB7-CI-EGFP-RBG (GFP-control) from the University of Pennsylvania Vector Core. Stereotaxic surgery was conducted on mice anaesthetized with isoflurane and administered ketoprofen (5 mg/kg) for pain management. For chemogenetic experiments, CAV2-Cre viral vector was bilaterally infused into the medial AON (10° angle toward midline; AP: +2.90, ML: ±1.10, DV: −3.42) and lateral AON (no angle; AP: +3.20, ML: ±1.10, DV: −3.90) at a volume of 0.1-0.2 μL and hM4D into the intermediate hippocampus (no angle; AP: −2.70, ML: ±2.20, DV: −2.00) and ventral hippocampus (10° angle away from midline; AP: −2.92, ML: ±2.15, DV: −4.90) at a volume of 0.3-0.4 μL. The position immediate to the rhinal fissure was used as a landmark to delineate intermediate and ventral parts. For optogenetic experiments, ArchT or GFP-control was bilaterally infused into the intermediate or ventral hippocampus in a volume of 0.3-0.4 μL, and optical fibers (200 μm core diameter, 0.39 NA; Thorlabs, Newton, NJ, USA) threaded through 1.25 mm-wide zirconia ferrules (Thorlabs) were bilaterally implanted into the lateral or medial AON, respectively. For anterograde tracing experiments, ChR2-mCherry was infused into the ventral hippocampus and ChR2-YFP was infused into the contralateral intermediate hippocampus. All infusions were made by means of pressure ejection at a rate of 0.1 μL/min through a cannula connected by Tygon tubing to a 10 μL Hamilton syringe (Hamilton, Reno, NV). A 15 minute interval was allotted after each infusion to limit the viral spread. The atlas of the mouse brain by Paxinos and Franklin (2007) was used to determine coordinates for guiding the stereotaxic infusion of viral vectors.

### Drugs

Clozapine-*N*-Oxide (CNO) obtained from the NIH was dissolved in a solution of 10% DMSO and 0.9% saline. A dose of 5 mg/kg of CNO was used in all behavioural experiments using hM4D-mCherry and mCherry-only expressing animals. All CNO treatments were separated by a minimum of 72 hours.

### Apparatus for optogenetic experiments

Inhibition of hippocampal terminals at the AON was conducted by illumination with green light (532 nm, 12 mW) generated by a diode-pumped solid state laser (Laserglow, Toronto, ON, Canada). The laser was connected to a 1 × 2 optical commutator (Doric Lenses, Quebec, QC, Canada) which divided the light path into two arena patch cables attached to the implanted optical fibers.

### Experimental Design

Unless otherwise noted, all tests took place in a 50 cm × 25 cm × 20 cm plexiglass open-topped cage. Odours were presented mixed with woodchip bedding in 3 cm wide, 1 cm high aluminum cups. Multiple identical odour cups were used such that an animal never investigated the same cup twice. The odours used included nutmeg, vanillin, coriander, banana, garlic, cinnamon, thyme, almond, onion, curry, ginger, savory, cumin, dill, jasmine, coffee, oregano, sage, and rosemary. The odours presented and the order of their presentation between animals was pseudorandomized. For habituation, mice were given 15 minutes of exploration time for each unique context prior to initial exposure. Each exposure was 5 minutes in length and inter-trial intervals were 15 minutes. All tests were video-recorded at 60 fps using a NIKON D5200 equipped with a 30 mm lens. An additional overhead video was recorded using a Logitech webcam. All videos were subsequently scored blind to the treatment groups. Exploration was strictly defined as head up sniffing, directed towards and within 1 cm of the odour source. This definition excludes the use of the odour cup for sitting or as support during rearing.

### Behavioural protocols

#### Olfactory Spatial Memory Test

In this paradigm adapted from Eacott and Norman (2004), mice were tested for memory of odour location in context. The test chamber was altered to produce two distinct contexts. Zebra-patterned paper was used to line the walls of context A whereas context B had transparent walls surrounded by red plastic cups and bedding on the floor. Each animal underwent two encoding phases and one retrieval phase. During the first encoding phase, mice explored context A where two highly distinct odours were placed at opposite ends of the chamber (odour 1 on the left and odour 2 on the right). Next, mice were removed from the chamber and placed in a holding cage. The mice were then returned to the chamber, except it was now configured as context B and contained both odours in opposite positions (odour 2 on the left and odour 1 on the right). The animals were allowed to explore both odours in their new positions before being placed back into the holding cage. For the retrieval phase, the chamber was reconfigured as context A but now two copies of one odour were presented on both sides of the chamber. Time spent investigating the odour cups was measured. The novel configuration consists of the familiar odour in a novel position within the original context. The initial context and left/right position of the odour cups were pseudorandomized.

#### Olfactory Temporal Order Memory Test

This paradigm is based on similar tests used previously to measure memory for the temporal order of objects^20, 21^. Mice were tested in a transparent chamber with spatial cues kept constant throughout the session. Each animal underwent three encoding phases and one retrieval phase. In the first exposure, mice were placed in the chamber with two copies of one odour presented on opposite sides of the arena. After exploring both copies, the animal was removed from the chamber and placed in a holding cage. This process was repeated two more times using different odours each time. During the retrieval phase the animal was returned to the chamber, but this time earlier and recently explored odours were presented on opposite sides. In this case, the odour explored earlier is ‘more novel’ given its temporal distance compared to the recently explored odour. The left-right positions of the first and last odours during the test were pseudorandomized. Time spent investigating both odours was measured.

#### Novel Odour Recognition Test

This test was given 72 hours after examining performance on temporal order memory and followed a similar paradigm. The animals underwent three encoding phases where two copies of a unique odour were presented in each. On the retrieval phase, mice were presented with the initially encountered odour and a previously unexplored odour on opposite sides of the chamber. Time spent investigating each odour was measured.

#### Olfactory Episodic Memory Test

Animals were first habituated to the apparatus which consisted of a 50 cm × 50 cm × 20 cm transparent plexiglass open field for a 30 minute period. The animals were then exposed to two encoding phases and one retrieval phase each separated by a one hour delay. In the first encoding phase, animals were given 10 minutes to explore two different odours located at two adjacent corners of the arena. In the second encoding phase, the animals were given an additional 10 minutes to explore another set of unique odours presented on the opposite adjacent corners. During the retrieval phase, all four odours were presented with the spatial position of one odour from each set exchanged. This presentation results in each odour possessing a unique spatiotemporal configuration-novel location and temporally distant (NL/TD), familiar location and temporally distant (FL/TD), novel location and temporally recent (NL/TR), and familiar location and temporally recent (FL/TR). Successful memory for an integrated (what, when, and where) memory results in a pattern of exploration such that the odour with the NL/TD configuration is preferentially investigated the most while the FL/TR configuration is investigated the least. Time spent investigating all four odours was measured for five minutes in the retrieval phase. Overhead videos were analyzed using the ANY-maze software to produce average heat maps of each treatment group’s position within the arena.

#### Context-Driven Odour Recall Test

In this paradigm adapted from Mandairon et al. (2014), mice were trained to associate a visually distinct context with an odour and subsequently tested for recollection of the odour when exposed to the context alone. The testing apparatus consists of a 50 cm × 30 cm × 20 cm plexiglas cage with colourful visual patterns pasted on the outside of the walls. A wooden applicator with a cotton swab tip was positioned 3 cm from the floor and 5 cm from one end of the chamber. Before introducing mice into the chamber, 100 μL of a pure odourant was applied to the cotton tip. Each mouse was randomly assigned a monomolecular odourant to be trained with among limonene, isoamyl acetate, nonane, and 1-pentanol. Mice were allowed to explore the context and the odorized cotton swab for 30 minutes per day for 9 consecutive days. On Day 10, mice were once again placed into the cage, however no odour was added to the cotton swab. Investigation time of the cotton swab was measured on day 9 and 10 for the first 5 minutes of their exposure to the context. Upon failure to detect the expected odour, mice behaviourally expressed memory by spending a greater amount of time investigating the cotton swab compared to their investigation time when the odour was present on Day 9.

### Histology

After behavioural testing, mice were transcardially perfused with Phosphate-Buffered Saline (PBS, pH 7.4), followed by 4% paraformaldehyde in phosphate buffer. Brain tissue was extracted and postfixed overnight at 4°C. The brains were then cryoprotected using a 30% sucrose in PBS solution. Coronal 40 μm thick sections were collected using a cryostat (Leica, Germany). The sections were slide-mounted, counterstained with 4′,6-diamidino-2-phenylindole 135 (DAPI) for five minutes, and subsequently coverslipped with Aquamount (Polysciences Inc, Warrington, PA). Wide-field fluorescent images were captured using a 4X objective lens on a fluorescent microscope (Olympus, Japan). Confocal images were captured using a 20X and 60X objective through a Quorum spinning disk confocal microscope (Zeiss). Adobe Photoshop CS6 (Adobe Systems Incorporated, San Jose, CA) was used to adjust the brightness and contrast of representative sections.

### Calculations and Statistical Analysis

The discrimination ratios were derived from the exploration time of odor-context pairings during the retrieval phase of each test. For all tests based on the spontaneous novelty preference paradigm, the discrimination ratio was calculated as the difference between the times spent exploring the novel and familiar odour-context configurations divided by the total amount of time investigating both odours. Here, a value of zero indicates that the animal investigated both odours to an equal extent. Positive values up to one indicate preference for the novel combination, whereas a negative value indicates greater investigation of the familiar odour-context pair.

For the episodic memory test, percent investigation time was calculated by dividing the amount of time spent investigating an individual odour-spatiotemporal configuration by the time investigating all odours and multiplied by 100%:

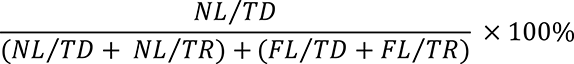

For analyzing spatial memory, the difference in the amount of time spent investigating odours with novel and familiar positions was divided by the total investigation time:

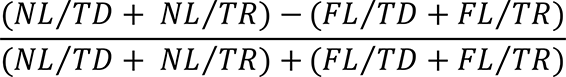

For temporal memory, the difference in the time spent investigating odours experienced earlier and later was divided by the total investigation time:

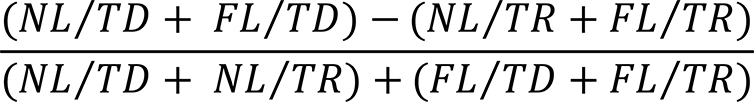

Performance in novelty preference-based paradigms was compared using t-test in hM4D experiments and a one-way ANOVA with testing group as the factor in optogenetic experiments. Percent investigation time data collected in the episodic memory test was analyzed using a two-way ANOVA with testing group and odour spatiotemporal configuration as factors. A two-way ANOVA was used to analyze data in the contextually-cued odour recall test for hM4D and optogenetic experiments, respectively, using experimental day and treatment group as factors. Where appropriate, Tukey’s multiple comparisons test was used for *post hoc* comparisons. Significance was defined as \**P*<0.05, \*\**P*<0.005, \*\*\**P*<0.0005, \*\*\*\**P*<0.0001.

## Acknowledgements

We thank Drs. Rutsuko Ito, Boyer Winters, Kaori Takehara, Paul Whissell, and John Yeomans for their valuable input regarding the collection and interpretation of data. This research was funded by operating grants to J.C.K. from the Canadian Institutes for Health Research (CIHR) (MOP 496401) and the Natural Sciences and Engineering Council of Canada (NSERC) (MOP 491009).

## Author contribution

A.J.A. and J.C.K. carried out the study conceptualization and experimental design. A.J.A performed and analyzed behavioural experiments. A.J.A and J.C.K wrote the manuscript.

## References

1. Tulving, E. Elements of episodic memory (Oxford Univ. Press, Oxford, 1985).

2. Tulving, E. & Markowitsch, H.J. Hippocampus 8, 198-204 (1998).

3. Eichenbaum, H. Annu Rev Psychol 68, 19-45 (2017).

4. Moser, E.I., Kropff, E., and Moser, M.-B. Annu Rev Neurosci 31, 69-89 (2008).

5. Eichenbaum, H. Nat Rev Neurosci 15, 732-744 (2014).

6. Gottried, J.A., Smith, A.P., Rugg, M.D., & Dolan, R.J. Neuron 42, 687-695 (2004).

7. Mandairon et al. Front Behav Neurosci 8, 138 (2014).

8. Tanaka et al. Neuron 84, 347-354 (2014).

9. Nyberg, L., Habib, R., McIntosh, A.R., & Tulving, E. PNAS 97, 11120-11124 (2000).

10. Rowe, T.B., Macrini, T.E., & Luo, Z.X. Science 332, 955-957 (2011).

11. Swanson, L.W., & Köhler, C. JNeurosci 6, 3010-23 (1986).

12. Brunjes, P.C., Illig, K.R., & Meyer, E. Bain Res Rev 50, 305-335 (2005).

13. Carson, K.A. Brain Res Bull 12, 629-634 (1984).

14. Aqrabawi et al. Nat Commun 7, 13721 (2016).

15. Ennaceur, A. & Delacour, J. Behav Brain Res 31, 47-59 (1988).

16. Barker et al. Nat Neurosci 20, 242-250 (2017).

17. Dere, E., Huston, J.P., De Souza Silva, M.A. Brain Res Protoc 16, 10-19 (2005).

18. Wheeler, M.E., Petersen, S.E., & Buckner, R.L. PNAS 97, 11125-11129 (2000).

19. Eacott, M.J., & Norman, G. J Neurosci 24, 1948-1953 (2004).

20. Hunsaker, M.R., Fieldsted, P.M., Rosenberg, J.S., & Kesner, R.P. Behav Neurosci 122, 643-650 (2008).

21. Mitchell, J.B., & Laiacona, J. Behav Brain Res 97, 107-113 (1998).

